# Differential LncRNA expression profile in the plasma of preeclampsia and normal pregnancies

**DOI:** 10.64898/2026.09.01.748658

**Authors:** Li Peng, Weisi Lai, Jian Huang, Yan Zhong

## Abstract

**Background:** Early detection of preeclampsia with noninvasive and reliable biomarkers is the foremost step for minimizing adverse effects during pregnancy. However, none maternal serum analyte evaluated so far is sufficiently accurate to allow recommending their routine use.

**Methods:** Microarray was used to first identify differentially expressed LncRNA and mRNA . Gene ontology (GO) and the Kyoto Encyclopedia of Genes and Genomes (KEGG) pathway analyses were performed to determine bio-functions and signaling pathways. Quantitative real-time polymerase chain reaction was used to validate the results of microarray. Finally, the lncRNA-mRNA co-expression network was constructed to find the interaction of genes.

**Results:** A total of 1476 differentially expressed lncRNAs and 594 mRNAs were identified. Both RNA-seq and RT-qPCR showed the expression of T241171,T338586, uc002ywy.3 was up-regulated, and the expression of ENST00000524858, T131416, T357032, uc.335+ was down regulated in the plasma of patients with preeclampsia.Pathway analysis showed the targeted mRNAs were enriched in apoptosis, sulfur metabolism, starch and sucrose metabolism. ceRNA network found that LncRNA-uc002ywy.3 might be the upstream regulator of miRNA-4498. LncRNA-uc002ywy.3/miRNA-4498 was predicted to interplay with genes involved in programmed cell death-1and its ligand (PD-1/PD-L1) pathway by KEGG analysis.

**Conclusions:** PD-1/PD-L1 signaling pathway may be involved in the development of preeclampsia. The dysregulated LncRNA-uc002ywy.3/miRNA-4498 shed light on a new layer involved in the regulatory network of preeclampsia.

## Introduction

Preeclampsia is major contributor to perinatal mortality and morbidity. It not only affects the outcome of pregnancy but also the long-term health of the mother and children. To date, there are no effective treatment for preeclampsia, other than termination of the pregnancy. Early detection and early intervention is proposed promising to minimize adverse effects of preeclampsia (1). However, none analyte for early dection so far is sufficiently accurate for clinical practice. Thus, it is necessary to futher study the pathogenesis of preeclampsia and search for more accurate analyte.

ncRNA is a type of functional RNA molecule that is not usually translated into protein.The regulatory ncRNAs can be divided into three types—lncRNA(> 200 nucleotides (nt)), miRNA (< 200 nt), and circRNA (circular structure).There is increasing evidence that miRNAs, lncRNAs, and circRNAs are widely involved in the pathogenesis of preeclampsia (2). miRNAs can regulate gene expression by targeting mRNA. We used to find miRNA1304-5p, miRNA 320a and miRNA 5002-5p were upregulated, while miRNA 188-3p, miRNA 211-5p, miRNA 4432 and miRNA 4498 were significantly downregulated in the plasma of preeclampsia patients(3).Alternatively, lncRNAs can serve as “sponges” to adjust the availability of miRNAs. Studies have shown that lncRNAs play important role in cell cycle regulation, immunity regulation, tumor genesis and epigenetic control(4) . Due to the essential role in gene regulation, lncRNAs have also been identified as potential biomarkers (2, 5). Although numerous studies have confirmed the differential expression of lncRNAs in placental tissues, studies on LncRNAs in the peripheral blood remain scarce. More research is required to elucidate the role of LncRNAs in the development of preeclampsia. In this study, we examined lncRNA expression profiles in preeclampsia blood samples compared with matched control samples. The results might contribute to the pathogenesis of preeclampsia, and provide new blood biomarkers for the diagnosis or treatment of preeclampsia.

## Methods

### Patient Samples

This is a prospective study at the Second Xiangya Hospital with recruitment period from Jan 1^st^ 2024 to Dec 31^st^ 2024, Written informed consent was obtained from all patients, and the study was approved by the Institutional Review Board of The Second Xiangya Hospital(approve number Z0553-01). Eligible women consisted of singleton pregnancy between 28 and 32 gestational weeks. Seven cases of preeclampsia and seven matched controls were included in the study. Preeclampsia was clinically diagnosed. The control group was made up of normal pregnancy without any complication. Multiple pregnancy,chronic hypertension,chronic hypertension with superimposed preeclampsia, gestational hypertension,eclampsia were excluded. The blood sample from each subject was snapfrozen in liquid nitrogen immediately after collection. Detailed information of all cases in the study was summarized in Table 1.

**Table 1.** Detailed information of all cases in the study.

|  | Control<br>Mean | (n=7)<br>SD | Case<br>Mean | (n=7)<br>SD | p |
| --- | --- | --- | --- | --- | --- |
| Gestational age(weeks) | 30 | 2 | 30.0 | 1.7 |  |
| Age(years) | 32.9 | 6.9 | 32.4 | 5.3 |  |
| Gravidity | 2.7 | 1.3 | 2.6 | 1.7 |  |
| Parity | 0.3 | 0.5 | 0.6 | 0.5 |  |
| Systolic blood pressure(mmHg) | 112.4 | 11.2 | 159 | 16.2 | <0.001 |
| Diastolic blood pressure(mmHg) | 73.1 | 6.4 | 107.6 | 11.9 | <0.002 |
| C-reactive protein(mg/l) | 5.6 | 1.1 | 11.3 | 12.8 |  |
| Hemoglobin(g/l) | 109 | 9.8 | 114.4 | 12 |  |
| White blood cells($10^9/l$) | 11.8 | 3.3 | 10.5 | 2 | |
| Platelet($10^9/l$) | 193.1 | 31.5 | 212 | 63.1 | |
| Fasting blood glucose(mmol/l) | 4.4 | 0.8 | 4.9 | 1.5 |  |
| Aspartate aminotransferase (u/l) | 19.4 | 5.8 | 20.8 | 6.4 |  |
| Alanine aminotransferase (u/l) | 14.1 | 6.5 | 17.7 | 6.5 |  |
| Albumin(g/l) | 33.9 | 4.5 | 28 | 3.1 | <0.05 |
| Total bilirubin(umol/l) | 9 | 5.3 | 6.5 | 1.5 |  |
| Prothrombin time(second) | 11.9 | 1 | 12.1 | 1.1 |  |
| Urea (mmol/l) | 3.1 | 0.9 | 5.2 | 2.4 | <0.05 |
| Creatinine(umol/l) | 46.4 | 14.9 | 61.5 | 24.2 |  |
| Uric acid(umol/l) | 229.2 | 31.1 | 443.4 | 114.8 | <0.001 |
|  | n | percentage | n | percentage |  |
| Abnormal Doppler | 0 | 0% | 4 | 57.14% | <0.05 |
| Fetal growth restriction | 1 | 14.29 | 5 | 71.43 | <0.05 |
| Urine protein |  |  |  |  |  |
| 0 | 6 | 85.71 | 1 | 14.29 |  |
| 1+ | 1 | 14.29 | 0 | 0 |  |
| 2+ | 0 | 0 | 4 | 57.14 |  |
| 3+ | 0 | 0 | 2 | 28.57 | <0.05 |

### RNA extraction

Total RNA was extracted from snap-frozen blood sample using TRIzol reagent (Invitrogen, Carlsbad, CA, USA) according to the manufacturer’s protocol. RNA quantity and quality were measured by NanoDrop ND-1000. RNA integrity was assessed by standard denaturing agarose gel electrophoresis.

### Microarray analysis

Arraystar Human LncRNA Microarray V4.0 was designed for the global profiling of human LncRNAs and protein-coding transcripts. About 40,173 LncRNAs and 20,730 coding transcripts could be detected by third-generation LncRNA microarray.

### RNA labeling and array hybridization

Sample labeling and array hybridization were performed according to the Agilent One-Color Microarray-Based Gene Expression Analysis protocol (Agilent Technology) with minor modifications. Briefly, mRNA was purified from total RNA after removal of rRNA (mRNA-ONLY™ Eukaryotic mRNA Isolation Kit, Epicentre). Then, each sample was amplified and transcribed into fluorescent cRNA along the entire length of the transcripts without 3’ bias utilizing a random priming method (Arraystar Flash RNA Labeling Kit, Arraystar). The labeled cRNAs were purified by RNeasy Mini Kit (Qiagen). The concentration and specific activity of the labeled cRNAs (pmol Cy3/μg cRNA) were measured by NanoDrop ND-1000. 1 μg of each labeled cRNA was fragmented by adding 5 μl 10 × Blocking Agent and 1 μl of 25 × Fragmentation Buffer, then heated the mixture at 60°C for 30 min, finally 25 μl 2 × GE Hybridization buffer was added to dilute the labeled cRNA. 50 μl of hybridization solution was dispensed into the gasket slide and assembled to the LncRNA expression microarray slide. The slides were incubated for 17 hours at 65°C in an Agilent Hybridization Oven. The hybridized arrays were washed, fixed and scanned with using the Agilent DNA Microarray Scanner.

### Quantitative Real-Time PCR validation

The remaining portion of the samples for lncRNA microarray was used for quantitative real-time polymerase chain reaction (qRT-PCR) validation. SuperScript III Reverse Transcriptase (Invitrogen, Grand Island, NY, USA) was used to reversely transcribe total RNA into cDNA accord-ing to the manufacturer’s instructions. Quan-titative real-time polymerase chain reaction (qRT-PCR) (Arraystar) was performed using the Applied Biosystems ViiA 7 RT PCR System and 2 × PCR Master Mix. The reaction conditions were set as follows: incubation at 95°C for 10 min, followed by 40 cycles of 95°C for 10 s and 60°C for 1 min. The relative expression levels of lncRNAs were calculated using the 2−ΔΔCt method and were normalized by β-actin (R).The data represented the means of three experiments.

### Gene ontology and KEGG analysis

Pathway analysis (based on KEGG, http://www.genome.jp/kegg/.) was performed to explore the significant pathways of the differentially expressed genes. Gene ontology (GO) (http://www.geneontol-ogy.org.) analysis was performed to determine the biological roles as molecular function, biological process, and cellular component of the aberrantly expressed mRNAs. P<0.05 and fractional disappearance rate (FDR) < 0.05 were used as thresholds to define markedly enriched GO terms/pathways.

### Construction of the Coding-non-coding Gene Coexpression Network

This co-expression network (CNC network) was performed between the validated lncRNAs and their related mRNAs based on the correlation analysis. The Pearson correlation coefficients (PCCs) of lncRNA and mRNA correlation analysis not less than 0.9 were chose to construct the network using Cytoscape software (version 2.8.3, The Cytoscape Consortium, San Diego, CA, USA). The solid lines represented a positive correlation, and the dashed lines indicated a negative correlation. The same color nodes represented co-expressed genes with a similar capacity chart. The node size indicated the gene expression, where nodes with more expressed gene coexpression had a more extensive relationship with the gene.

### Statistical analysis

SPSS Statistics (version 13.0; SPSS Inc., Chicago, IL, USA) was used for statistical analysis. Results were expressed as mean±standard deviation (SD) . Differences between the preeclampsia group and the control group were analyzed using Student’s t-test. Spearman correlation analysis was used to detect the relationship between lncRNAs and mRNAs. A P value of <0.05 was regarded as statistically significant.

## Results

### Clinical Characteristics of the Study Population

Blood samples were collected from a total of 14 patients, including 7 non-preeclampsia healthy subjects and 7 patients with preeclampsia. Systolic (159 ± 16.2 mmHg versus 112.4 ± 11.2 mmHg) and diastolic blood pressure (107.6 ± 11.9 mmHg versus 73.1 ± 6.4 mmHg) were higher in the preeclampsia group than in the non-preeclampsia group. Further, there were statistically significant differences in urinary protein, fetal growth restriction, abnormal Doppler between the preeclampsia patients and the non-preeclampsia control subjects. Additionally, preeclampsia patients had obviously lower albumin level than non-preeclampsia patients did (28 ± 3.1versus 33.9 ± 4.5 g/l, respectively). There were no significant differences in age, C-reactive protein, hemoglobin, white blood cells, or fasting blood glucose between the preeclampsia and non-preeclampsia groups (Table 1).

### Overview of lncRNA Profiles

Based on the lncRNAs expression profiles, differentially expressed lncRNAs could be found between the preeclampsia (test) and normal samples (control) with a test/control of 0.37(Figure 1). Volcano plots were constructed to display the expression profile for all detected lncRNAs (Figure 2). We determined 1476 differentially expressed human lncRNAs (853 up regulated; 623 down regulated, p<0.05) (Table plus 1). The heat map was shown in Figure 3.

**Figure 1.**
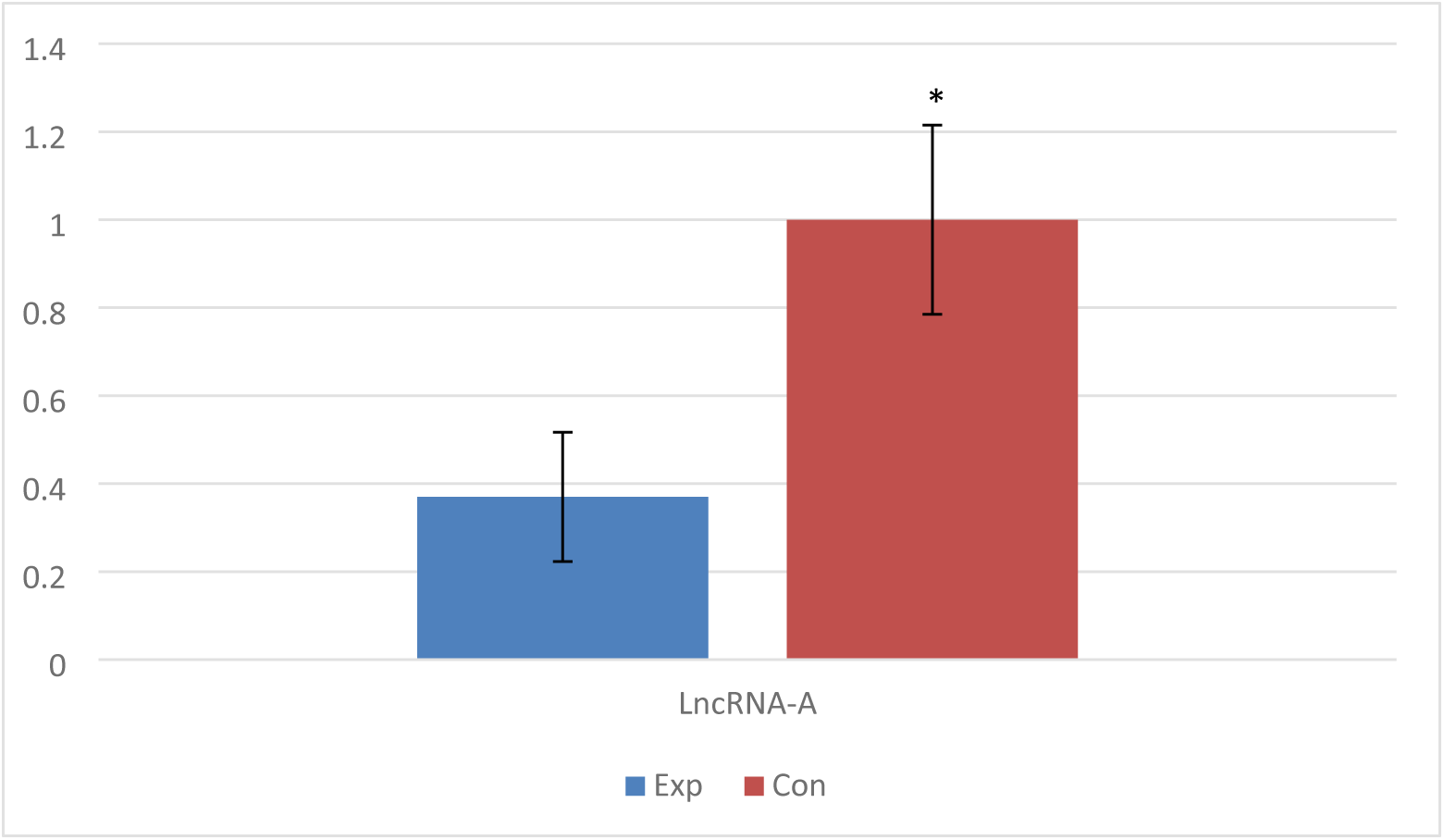
Differentially expressed lncRNAs between the preeclampsia (Exp) and normal samples (Con)

**Figure 2.**
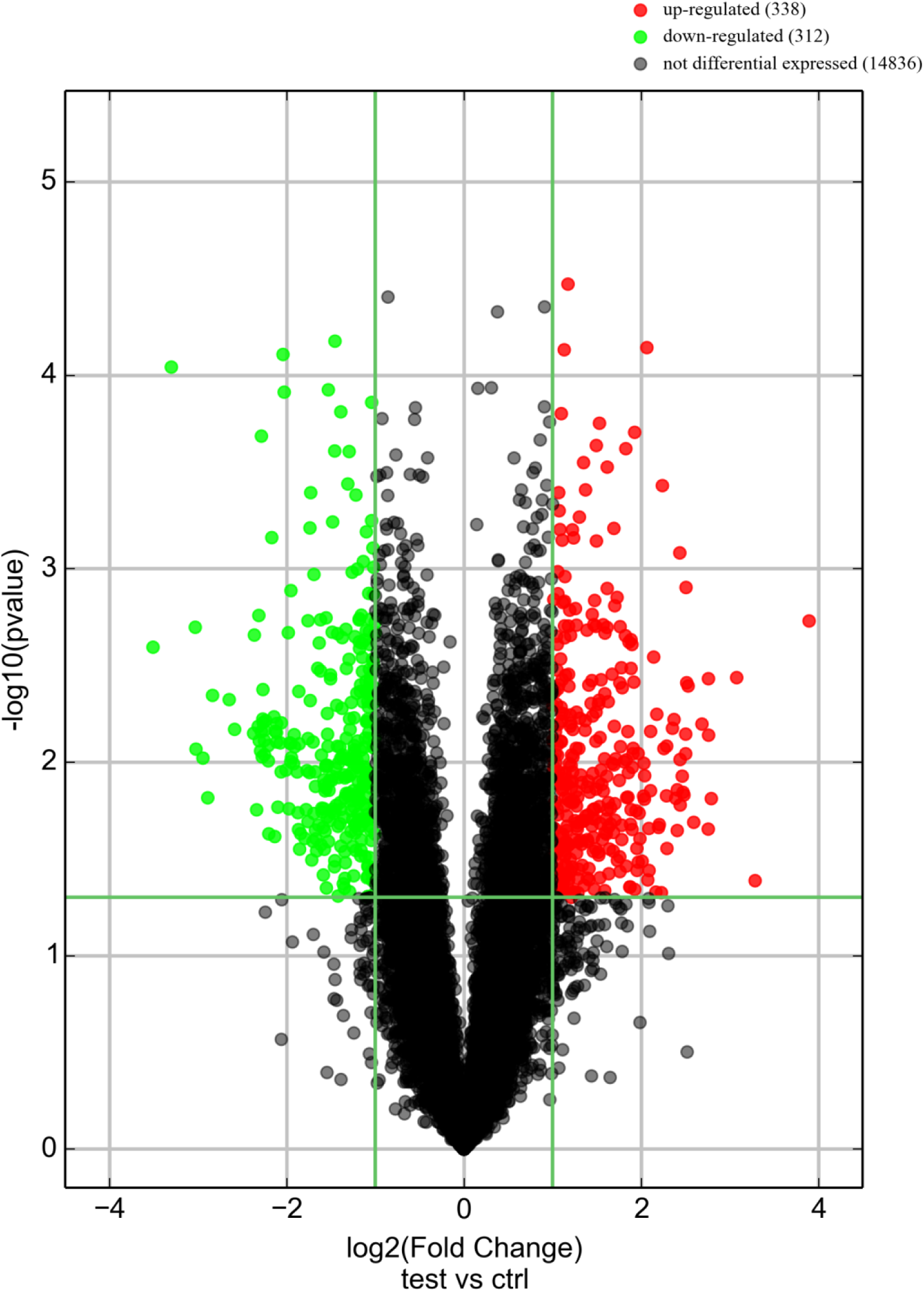
Volcano plots of the expression profile for all detected lncRNAs between the preeclampsia (test) and normal samples (Ctrl)

**Figure 3.**
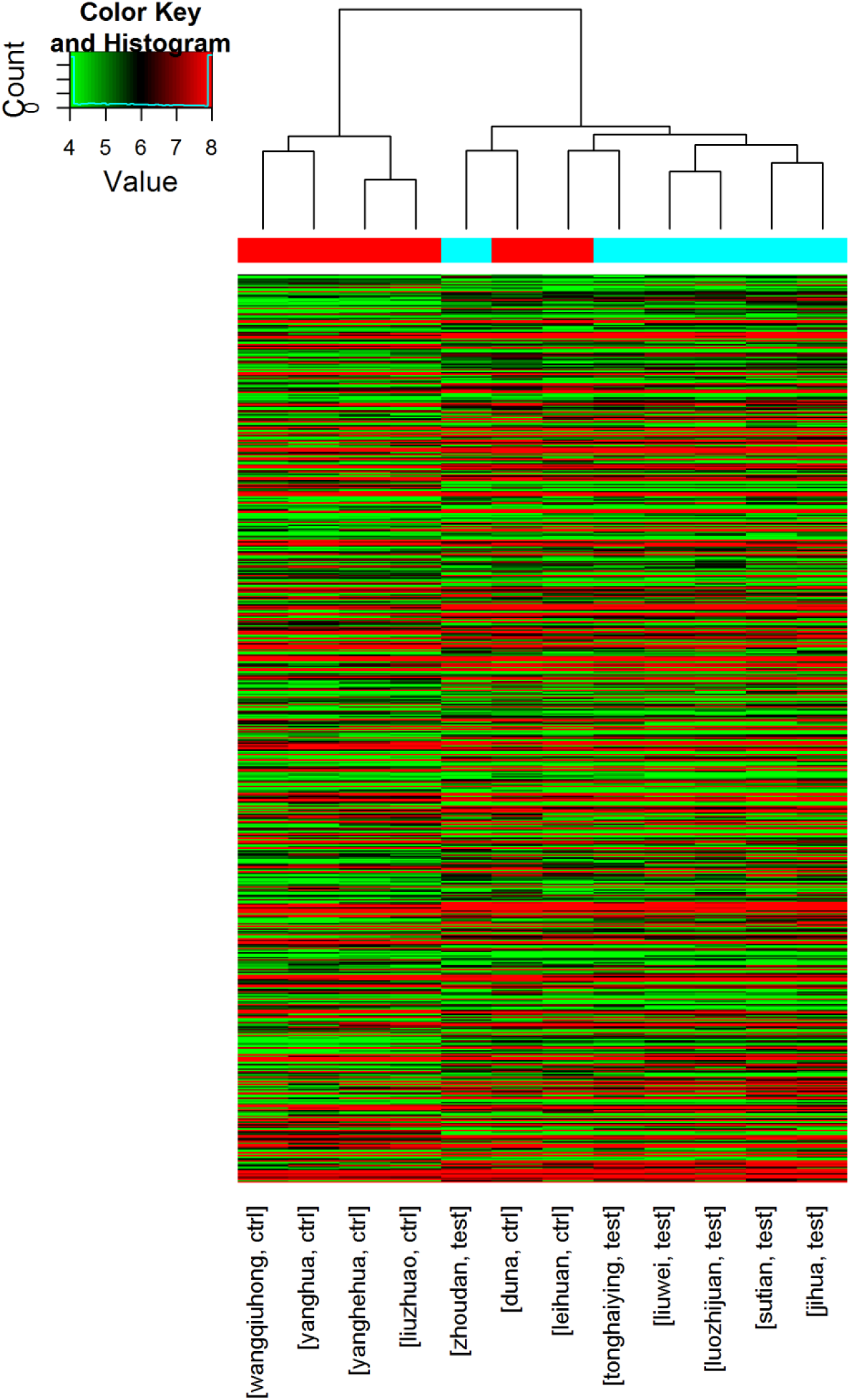
Heat map of lncRNAs expression profiles between the preeclampsia (test) and normal samples (Ctrl)

### Overview of mRNA Profiles

594 differentially expressed mRNAs were identified with a fold change cut-off of 2.0 (326 up-regulated; 268 down-regulated, p<0.05, Table plus 2). NM_001024630 and NM_207325 were the most up- and down regulated mRNA transcripts with the fold change of 13.06 and 12.6, respectively. The heat map was shown in Figure 4. The volcano plots displayed the expression profile for all detected mRNAs (Figure 5).

**Figure 4.**
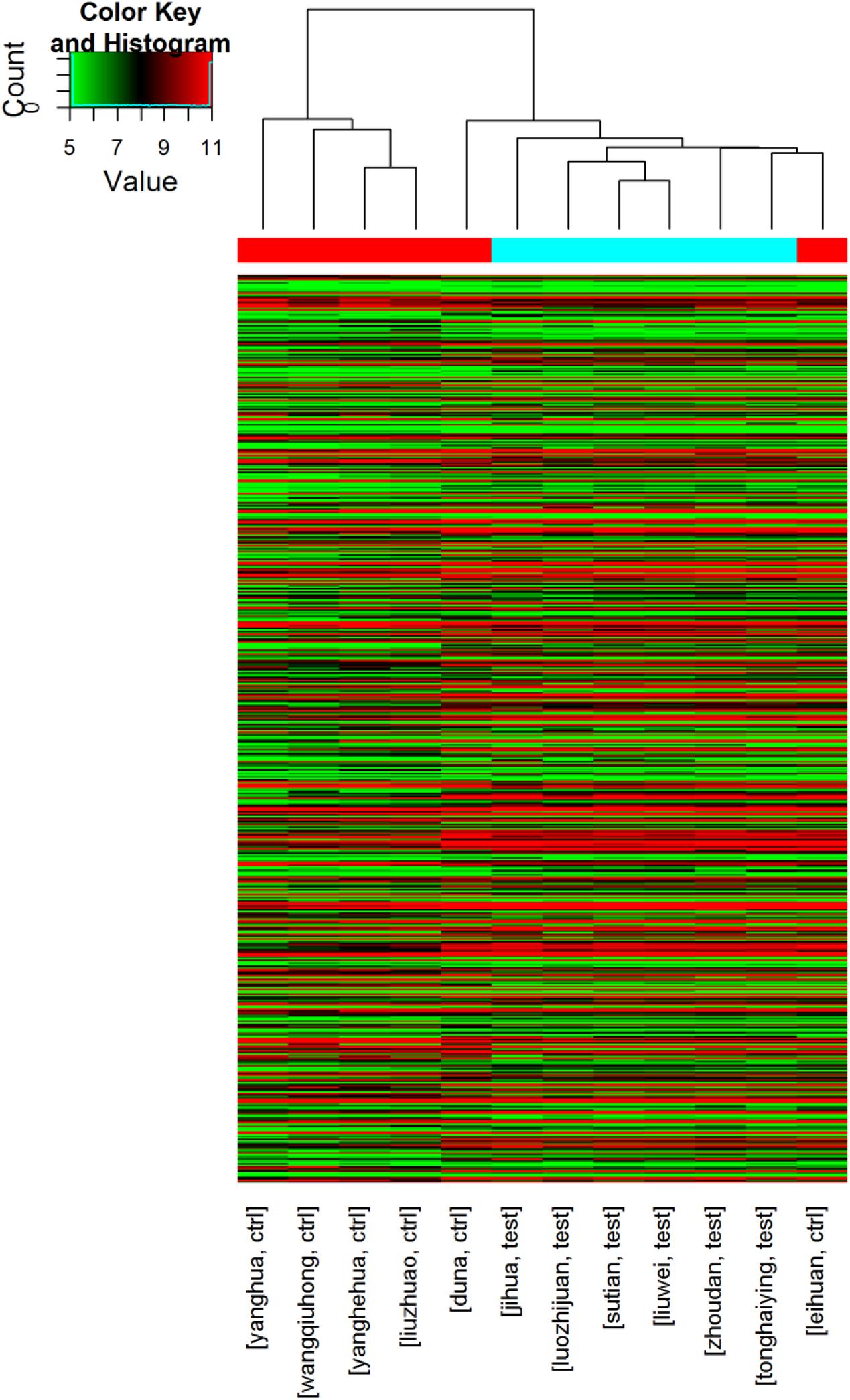
Heat map of mRNAs expression profiles between the preeclampsia (test) and normal samples (Ctrl)

**Figure 5.**
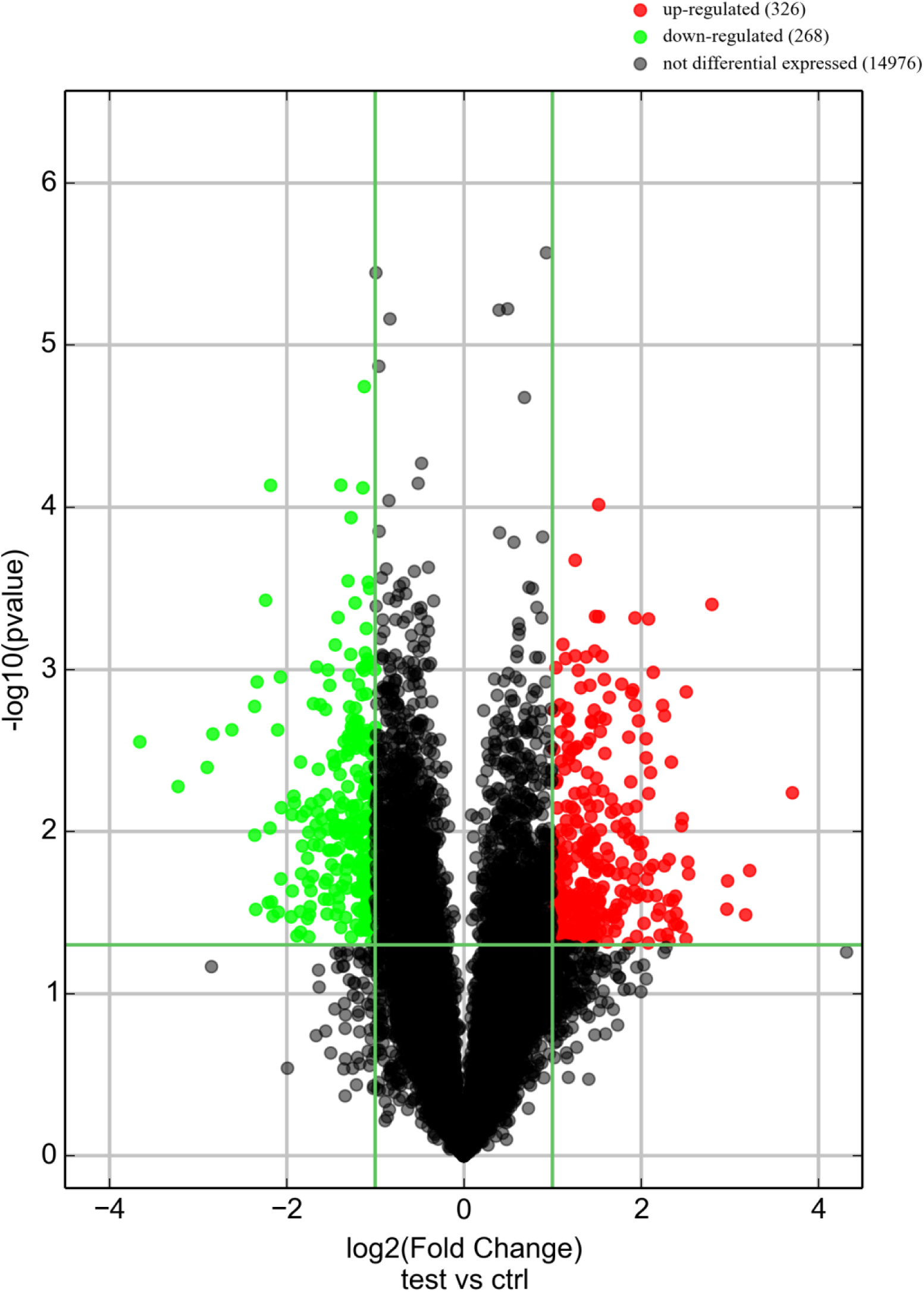
Volcano plots of the expression profile for all detected mRNAs between the preeclampsia (test) and normal samples (Ctrl)

### GO analyses

GO analyses classified the differentially expressed mRNAs to three categories: biological process (BP), molecular function (MF), cellular component(CC). The BP terms of up regulated mRNAs included immune response, defense response, immune system process, etc. The MF terms of up regulated mRNAs included chemokine activity, RAGE receptor binding, IgG binding, etc(Figure 6).The CC terms of up regulated mRNAs included extracellular space,bounding membrane of organelle,whole membrane, etc. The BP terms of down regulated mRNAs included regulation of G1/S transition of mitotic cell cycle, regulation of cell cycle G1/S phase transition,G1/S transition of mitotic cell cycle, etc (Figure 7). The MF terms of down regulated mRNAs included 2 iron, 2 sulfur cluster binding, thiol-dependent ubiquitin-specific protease activity, ubiquitinyl hydrolase activity, etc. The CC terms of down regulated mRNAs included lamellar body, transcription factor complex, Cul4-RING E3 ubiquitin ligase complex, etc.

**Figure 6.**
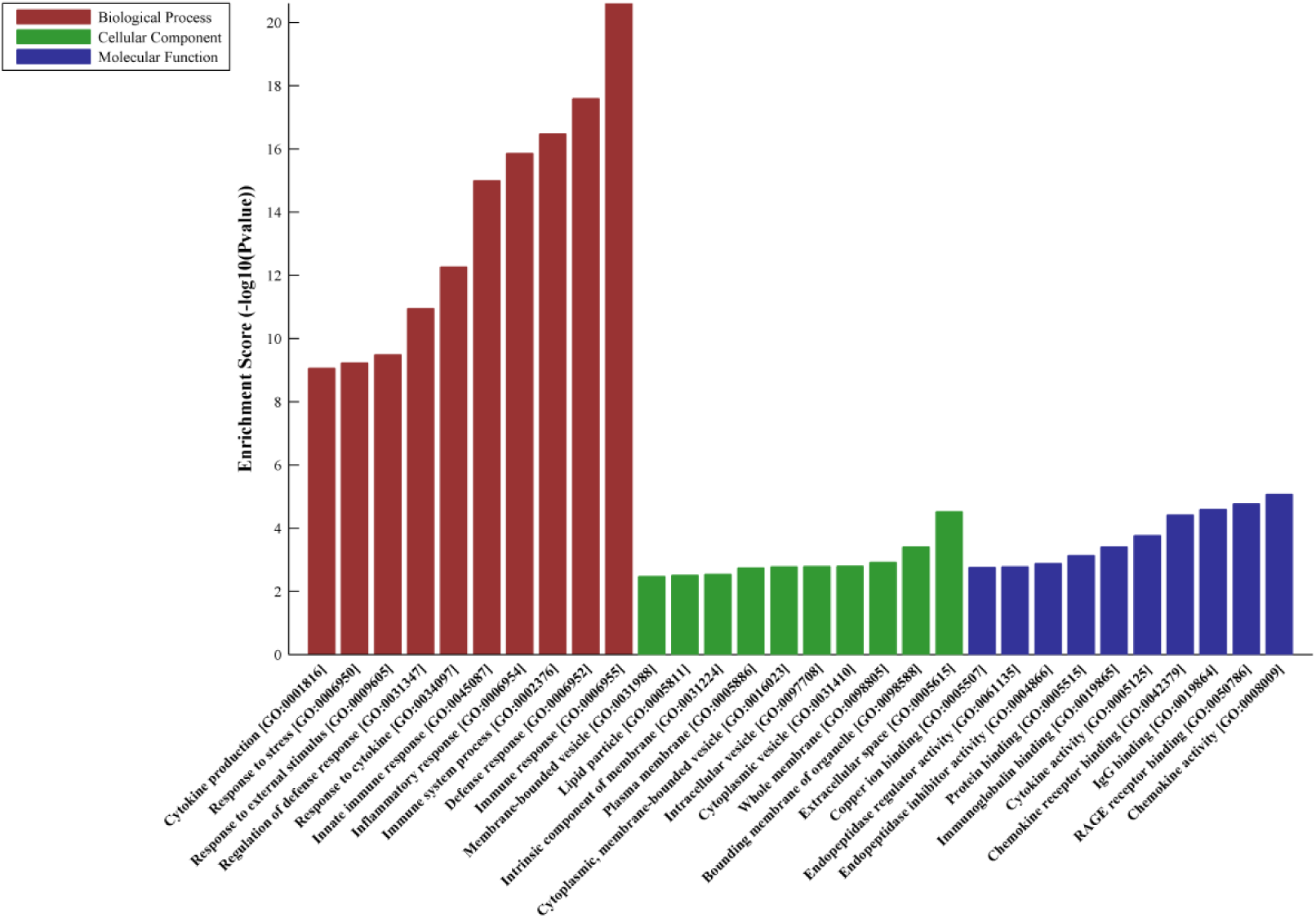
GO analyses of up regulated mRNAs

**Figure 7.**
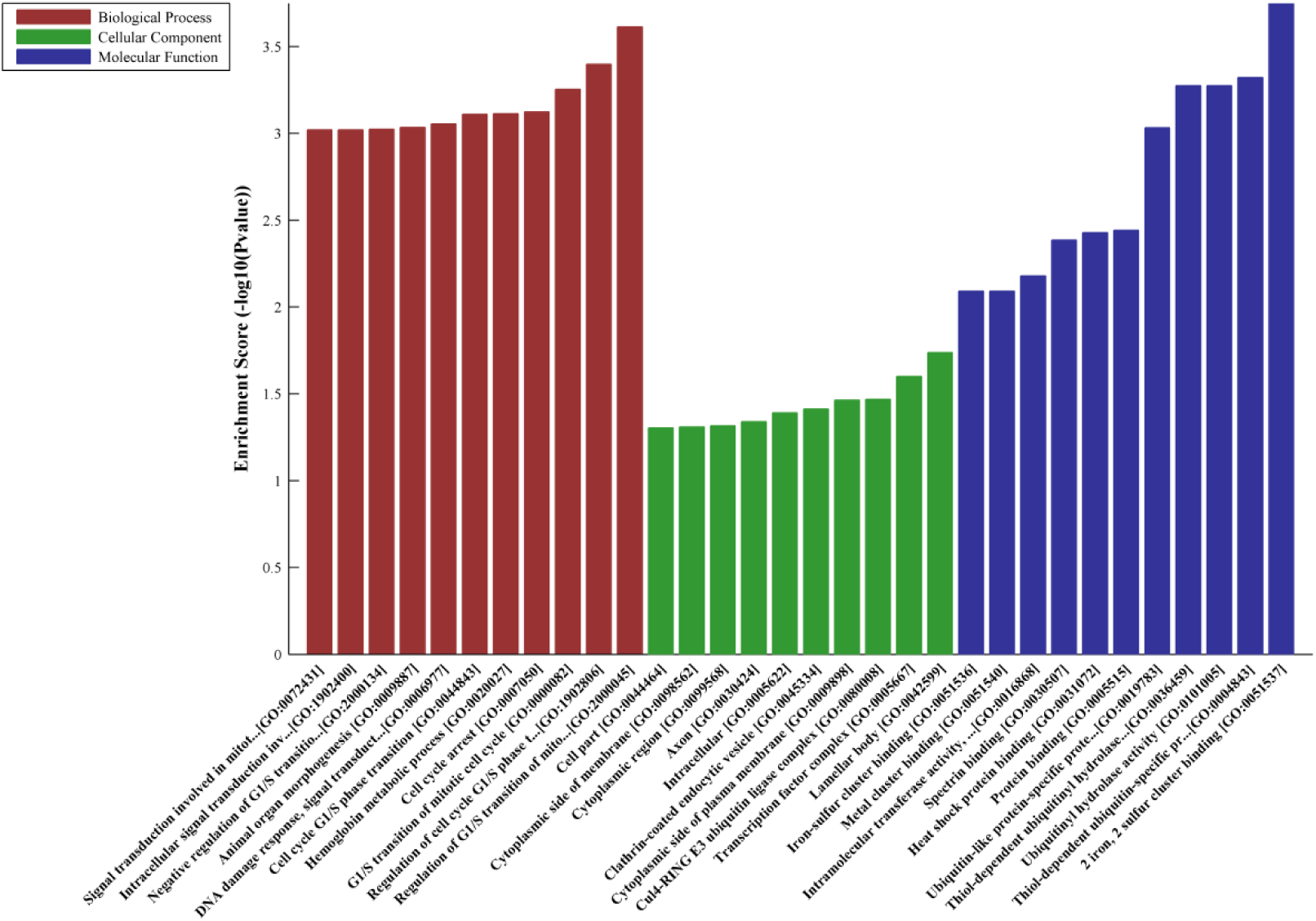
GO analyses of down regulated mRNAs Signaling pathway analysis

Signaling pathway analysis showed there were 28 pathways for up regulated mRNAs, involving toll-like receptor signaling pathway, NF-kappa B signaling pathway (Figure 8). There were 7 pathways for down regulated mRNAs, involving thiamine metabolism, TGF-beta signaling pathway (Figure 9).

**Figure 8.**
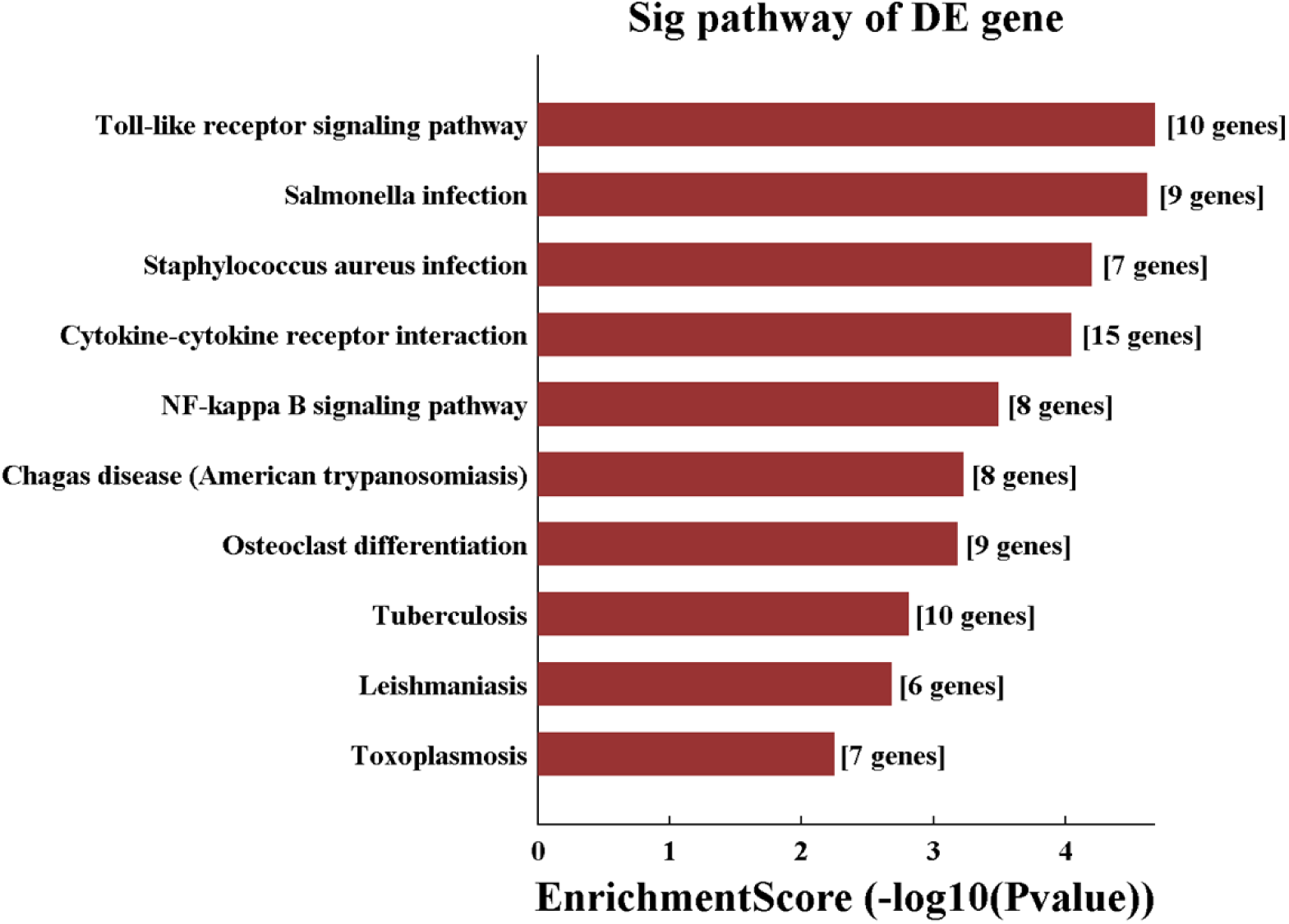
Pathway analysis for up regulated mRNAs

**Figure 9.**
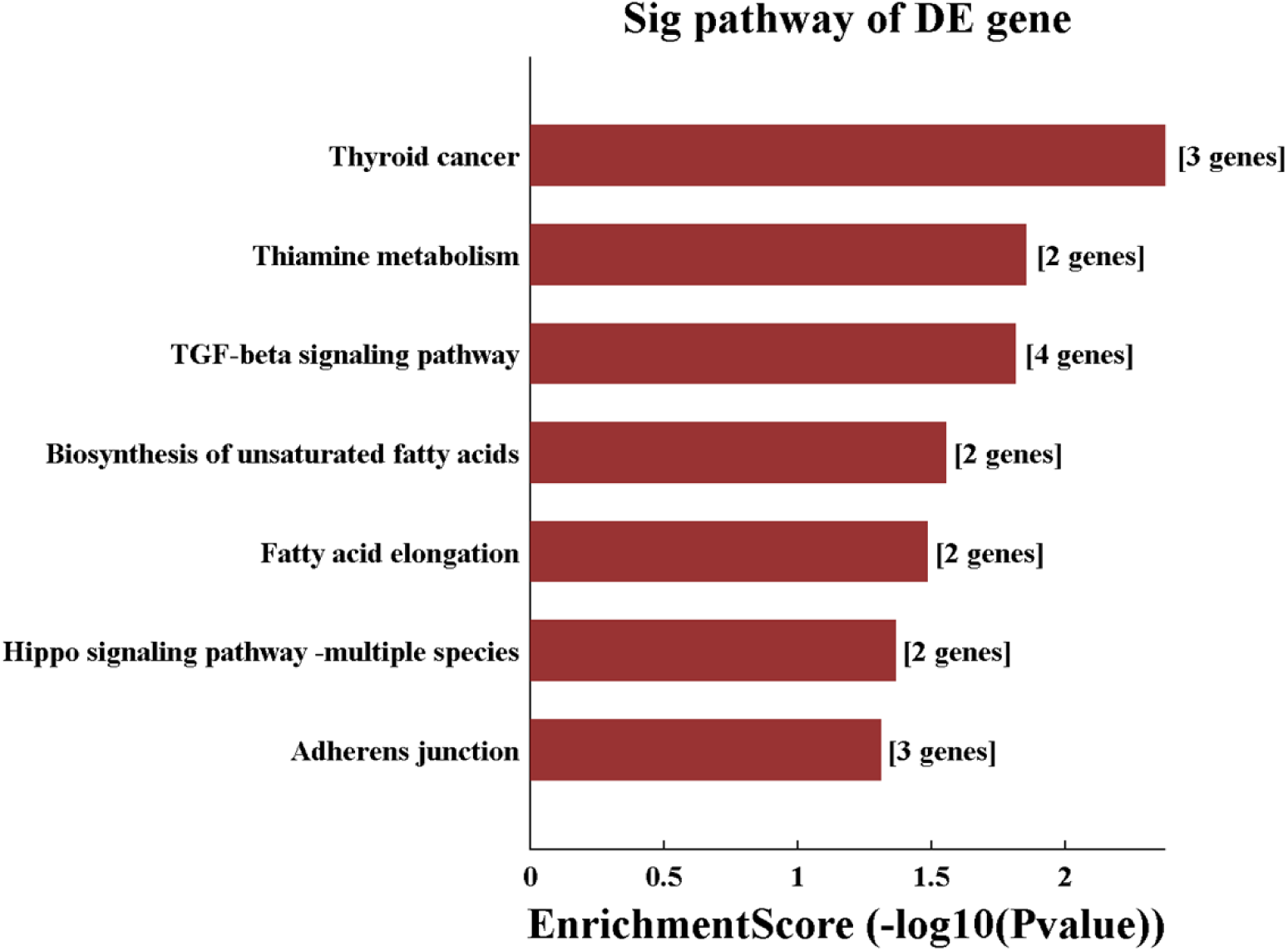
Pathway analysis for down regulated mRNAs

### Real-time Quantitative PCR Validation

Three most significant up-regulated lncRNAs(T241171,T338586, uc002ywy.3) and four most down-regulated lncRNAs(ENST00000524858, T131416, T357032, uc.335+) were selected to validate the expression changes detected by RNA-seq, . Their expression was examined using RT-qPCR. Both RNA-seq and RT-qPCR showed that the expression of T241171, T338586, uc002ywy.3 was up-regulated, and the expression of ENST00000524858, T131416, T357032, uc.335+ was down regulated (Figure 10).

**Figure 10.**
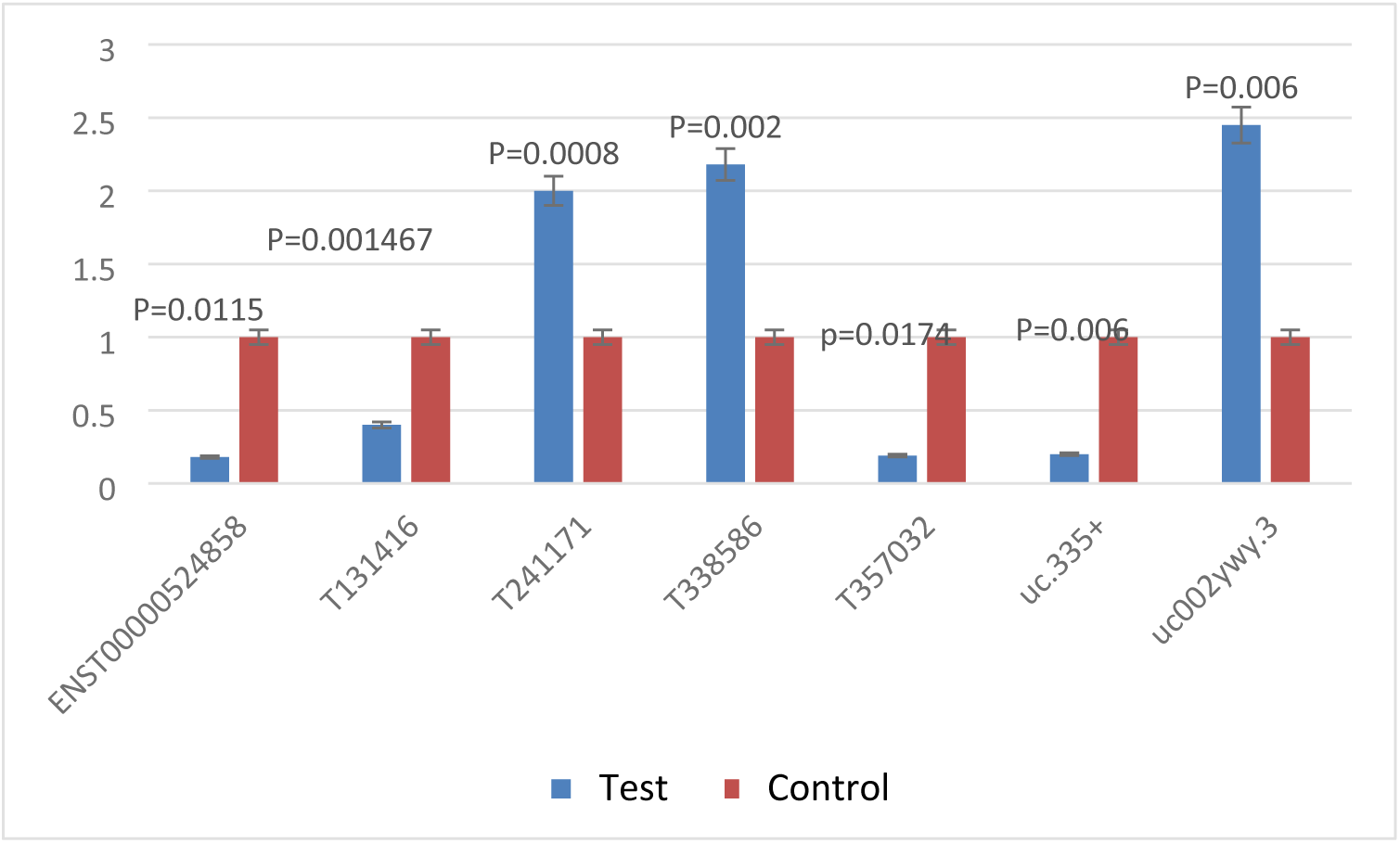
Real-time Quantitative PCR Validation lncRNAs expression

### Construction of the Coding-non-coding Gene Coexpression Network

We constructed the lncRNA and mRNA co-expression network profiles based on seven validated differentially expressed lncRNAs and interacted mRNAs (Figure 11). We further performed GO and pathway analysis based on the results of the co-expression network. GO analysis showed the BP terms of targeted mRNAs included neutrophil activation, myeloid leukocyte activation, granulocyte activation, etc(Figure 12). The MF terms included enzyme inhibitor activity, RAGE receptor binding, IgG binding, etc. The CC terms included secretory granule, secretory vesicle, cytoplasmic vesicle, etc. Pathway analysis showed that the targeted mRNAs were enriched in apoptosis, sulfur metabolism, starch and sucrose metabolism (Figure 13). Through ceRNA network, we found that the LncRNA-uc002ywy.3 might be the upstream regulator of miRNA-4498 (Figure 14). In addition, LncRNA-uc002ywy.3/miRNA-4498 was predicted to interplay with genes involved in programmed cell death-1and its ligand (PD-1/PD-L1) pathway (pathway ID: rno04740) by KEGG analysis (Figure 15、16, Table plus 3), including EGFR、 MAPK1、MYD88、STAT3、NFATC2、PIK3R3、PTPN11、RAF1、TRAF6.

**Figure 11.**
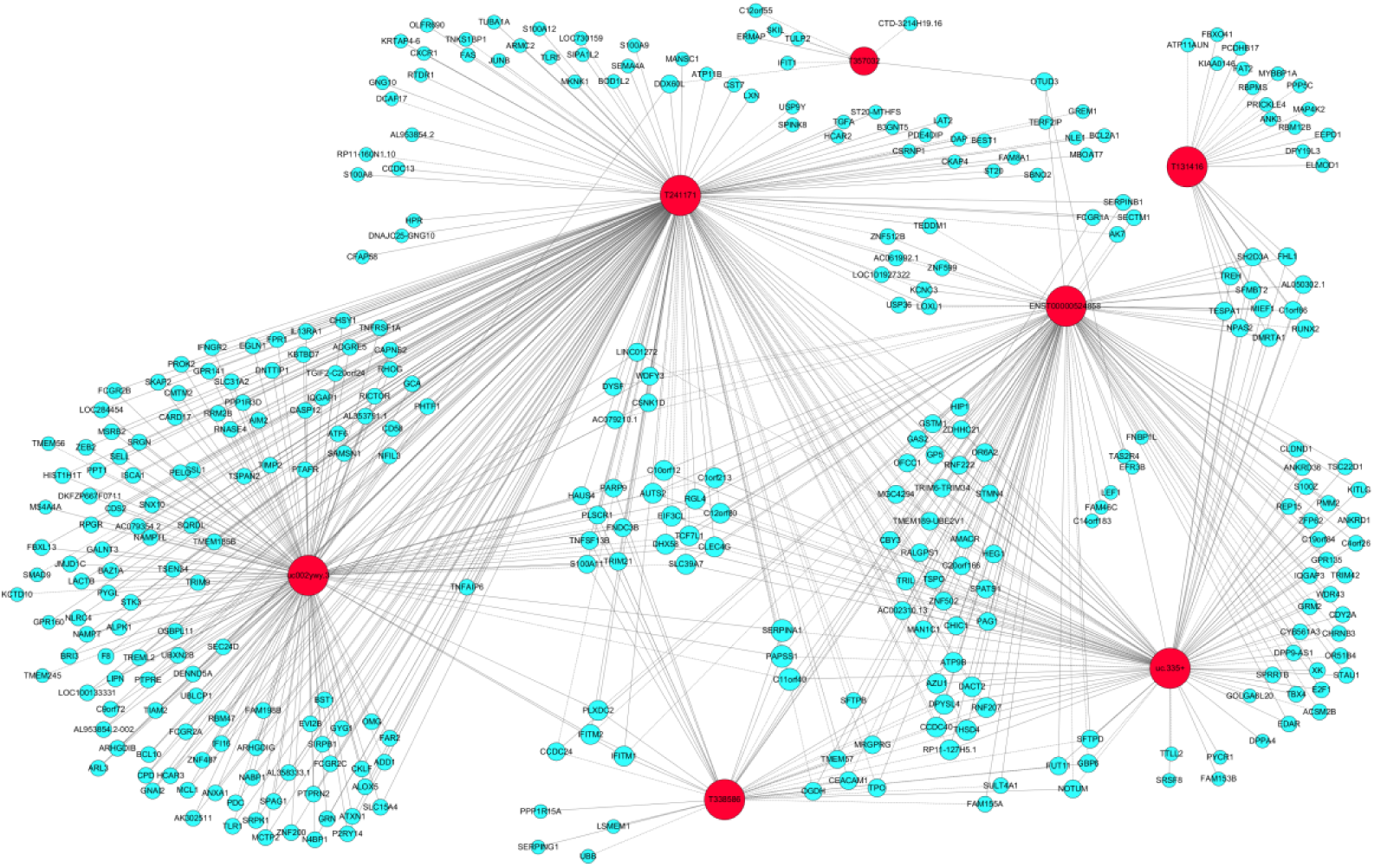
lncRNA and mRNA co-expression network profiles

**Figure 12.**
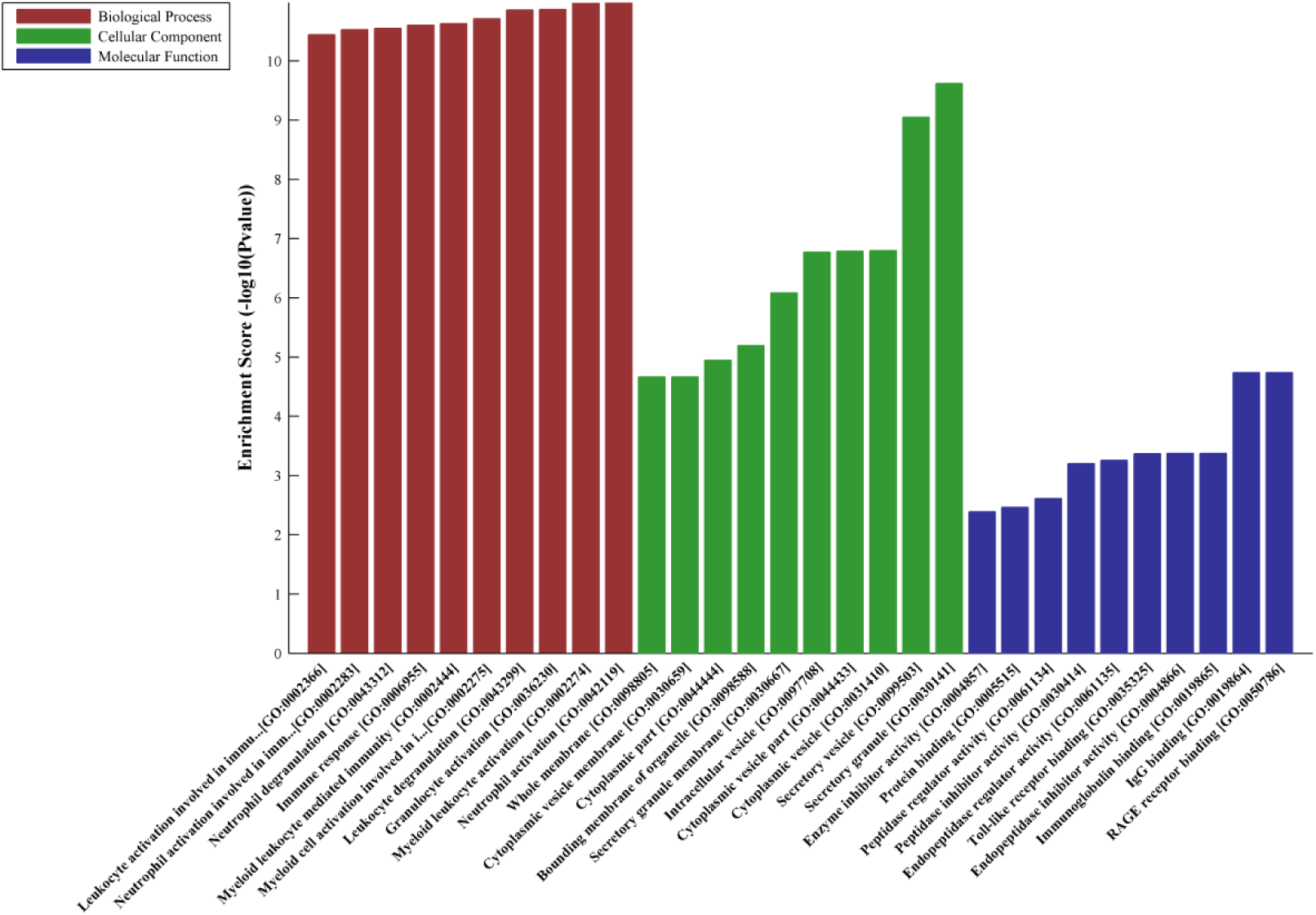
GO analysis of the co-expression network

**Figure 13.**
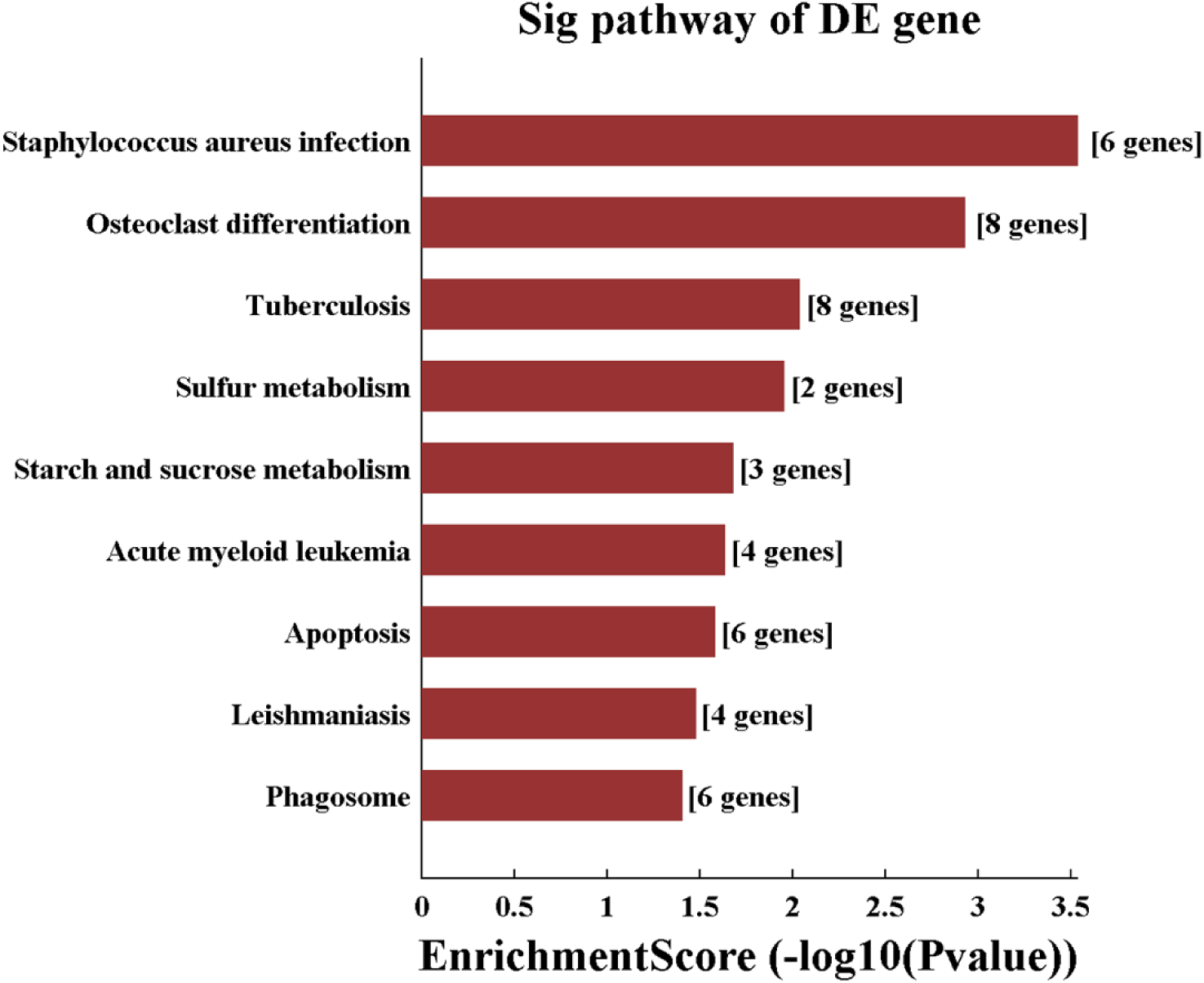
Pathway analysis of the co-expression network

**Figure 14.**
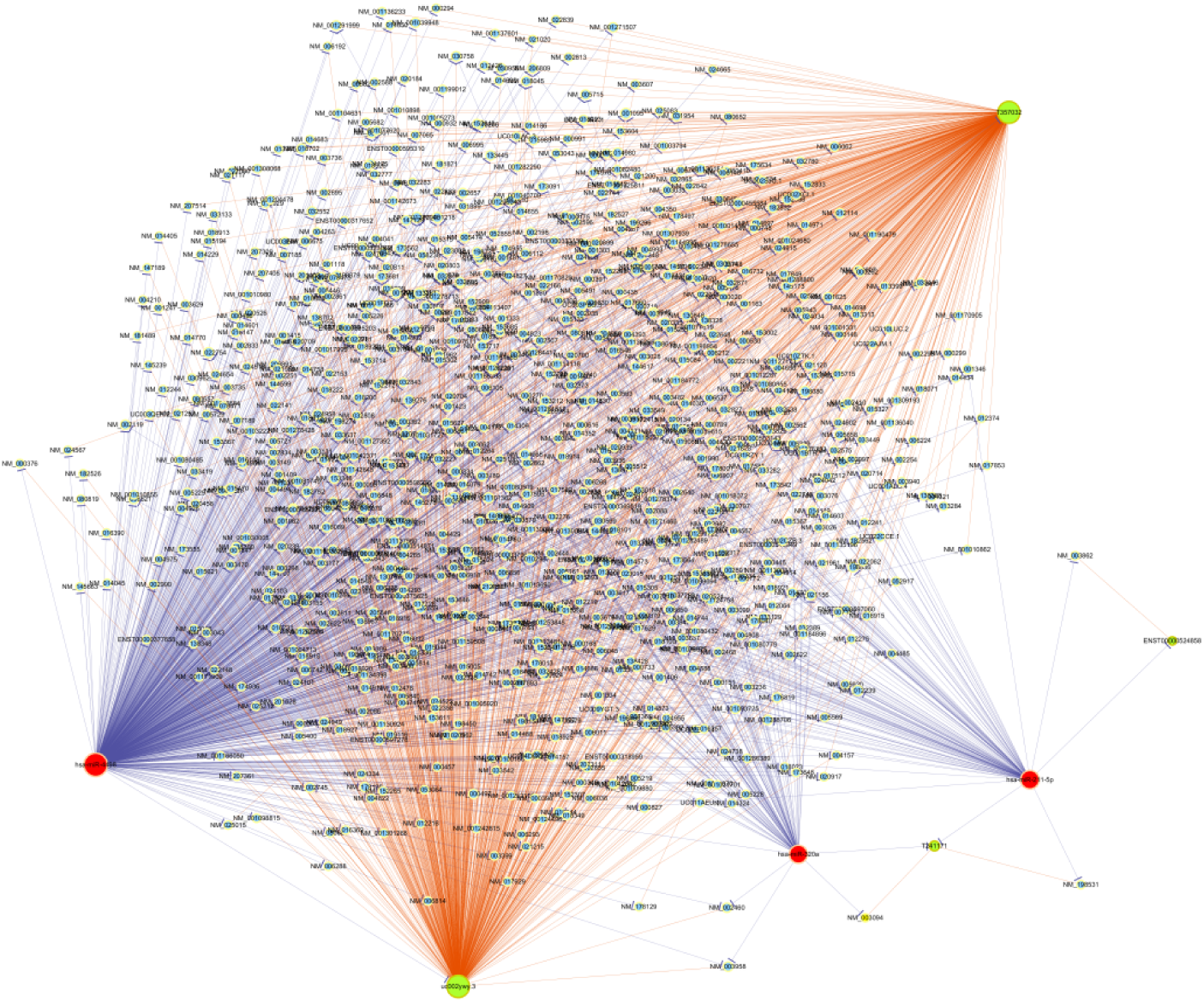
ceRNA network shows LncRNA-uc002ywy.3 might be the upstream regulator of miRNA-4498

**Figure 15.**
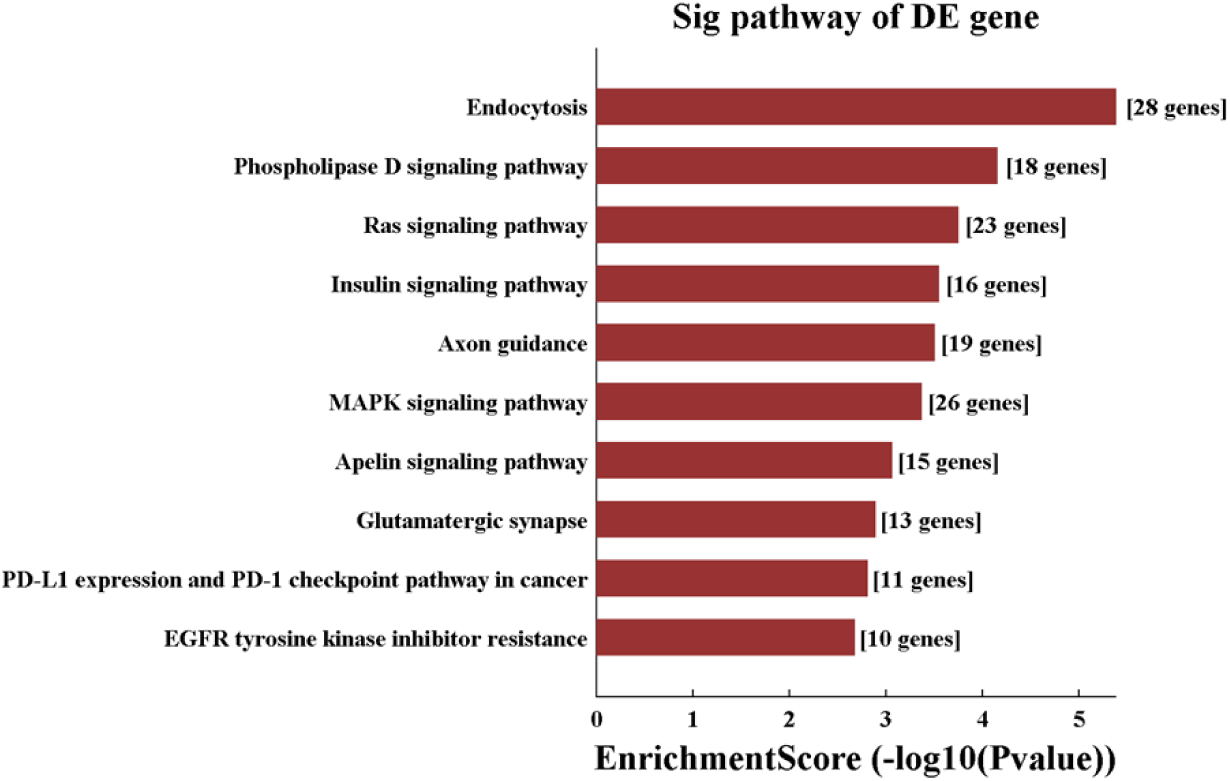
LncRNA-uc002ywy.3/miRNA-4498 was predicted to interplay with genes involved in PD-1/PD-L1 pathway

**Figure 16.**
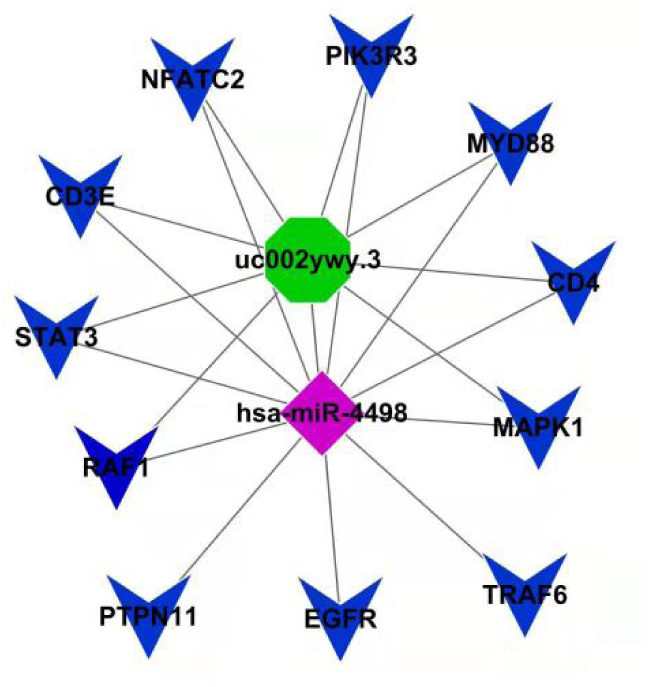
LncRNA-uc002ywy.3/miRNA-4498 was predicted to interplay with genes involved in PD-1/PD-L1 pathway

### Subgroup analysis

Combination analysis of antisense lncRNAs and mRNAs

Combination analysis of antisense lncRNAs and coding gene data showed 10 antisense lncRNAs and accordant mRNAs expressed differentially between preeclampsia and control group (Table plus 4).

Combination analysis of lincRNAs and nearby mRNAs

Combination analysis of lincRNAs and coding gene data showed 66 lincRNAs and nearby mRNAs expressed differentially between preeclampsia and control group (Table plus 5).

## Discussion

This study depicted the lncRNA and mRNA profile in serum for preeclampsia in the third semester. We identified LncRNA-uc002ywy.3 might be the upstream regulator of miRNA-4498. LncRNA-uc002ywy.3/miRNA-4498 was predicted to interplay with genes involved in PD-1/PD-L1 pathway.

LncRNAs were reported to involve in the occurrence and development of preeclampsia through impact on the biological functions of trophoblasts (including cell proliferation, migration, invasion, and apoptosis), immune regulation, epigenetic regulation, decidualization, and energy metabolism (5).The changes in the expression of lncRNAs in peripheral maternal blood during pregnancy had been proposed as possible biomarkers of preeclampsia. Wang et al. found the expression level of lncRNA NONHSAT116812 and NONHSAT145880 was significantly lower and higher, respectively, in term placenta of preclampsia women, strongly correlated with their expression on the plasma of the same women 48h before delivery(6).In our study, the expression of lncRNA T241171,T338586, uc002ywy.3 was up-regulated, and the expression of lncRNA ENST00000524858, T131416, T357032, uc.335+ was down-regulated in the third trimester peripheral maternal blood in preeclampsia patients. The difference of results between our and other study might be due to the different gestational age of sampling. Thus, gestational age should be taken into consideration when using lncRNA as biomarker for preeclampsia.

We found LncRNA(T241171, T338586,uc002ywy.3) associated with apoptosis and TNF were up regulated, whereas the down regulated genes participated in G1/S transition of mitotic cell cycle. The BP terms of Go analysis also showed a decrease in G1/S transition of mitotic cell cycle. This result was consistent with previous studies(4). It is reported that LncRNAs MALAT-1(7), MEG3(9) could induce cell arrest and apoptosis, while LncRNA GASAL1could promote trophoblastic proliferation and progression (8). All these indicating increased cell apoptosis and decreased cell proliferatioin was related to the pathophysiology in third trimester for reeclampsia. Some LncRNAs promoted this process while others could inhibit it. It might be a therapeutic target to treat preclampsia by inhibiting the LncRNAs promoting apoptosis and stimulating the LncRNAs promoting proliferation.

In our study, GO analysis showed the BP terms of targeted mRNAs included neutrophil activation, myeloid leukocyte activation, granulocyte activation, indicating chronic inflammation. Proper release of cytokines by the activated leukocytes or uterine epithelial cells is necessary for fetus implantation. While in preeclampsia, decreased placental perfusion and oxidative stress would increase the release of cytokines and antiangiogenic factors, leading to inflammation and endothelial dysfunction of numerous organs and systems (11). Thus, the BP terms of targeted mRNAs in our study correctly showed the characteristics of preeclampsia. Our pathway analysis showed the functions of altered genes were enriched in TLR signaling pathway and nuclear factor-kappa B (NF-κB) signaling pathway.

Uncontrolled activation of TLRs was suggested associated with pregnancy-related problems(10). TLRs could relay the signaling via the intracellular signaling adapter protein, the myeloid differentiation factor 88 (MyD88)-dependent pathway. The MyD88-dependent pathway leaded to the activation of early phase NF-κB, resulting in the production of pro-inflammatory cytokines, including IL-1β, IL-6, IL-12, and TNF-α(10). Oxidative stress in the preeclampsia women could elevate the activity of NF-κB signaling (12), and regulate epithelial-mesenchymal transition(EMT) processes(13) . Serum TLR4 and NF-κB p65 could be used as a biomarker for predicting cytokine environment and its influence on the immune cells (14). Both our study and previous study demenstrated the role of TLRs in the pathophysiology of preeclampsia. TLRs may be key mechanism mediating preeclampsia development and may provide novel biomarkers for preeclampsia.

Our study found the most upregulated genes participate in immune and defense response. LncRNA-uc002ywy.3 targeted miRNA4498 and LncRNA-uc002ywy.3/miRNA-4498 might influence the PD-1/PD-L1 signaling pathway. The interaction between PD-1 and PD-L1 was suggested associated with the Treg/Th17 imbalance in human pregnancy (15, 16). The imbalance of the Treg and Th17 response could be a result of TLR4 activation which created a pro-inflammatory environment leading to preeclampsia (17) . Previous studies found ncRNAs can regulate Treg/Th17 balance, such as miR-146a-5p(18) and LncRNA-NEAT1(19) .

LncRNA-NEAT1 was increased in patients with preeclampsia, knockdown of NEAT1 improved Treg/Th17 imbalance(20). Elevated miR-210 was found contribute to preeclampsia via inhibiting anti-inflammatory Th2-cytokines(21). Whether LncRNAs regulate Treg/Th17 imbalance via PD-1/PD-L1 signaling pathway requires to be further explored. How LncRNA-uc002ywy.3/miRNA-4498 participates in the development of preeclampsia via PD-1/PD-L1 signaling pathway would be the next step in our future study.

## Conclusion

In this study, we identified the differential expression patterns of lncRNAs and mRNAs in third trimester preeclampsia patients. Through the relevant analysis, we found that PD-1/PD-L1 signaling pathway may be involved in the development of preeclampsia. The dysregulated LncRNA-uc002ywy.3/miRNA-4498 shed light on a new layer involved in the regulatory network of preeclampsia. Further study is required to validate our result on the pathogenesis of LncRNA-uc002ywy.3/miRNA-4498 participating in the development of preeclampsia .

## List of abbreviations

BP: biological process
CC: cellular component
CNC: Coding-non-coding
GO: Gene ontology
KEGG: Kyoto Encyclopedia of Genes and Genomes
MF: molecular function
MyD88: myeloid differentiation factor 88
PD-1: programmed cell death-1
PD-L1: programmed cell death-1 ligand
TLR: toll-like receptor

## Declarations

### Ethics approval and consent to participate

The study was approved by the Institutional Review Board of The Second Xiangya Hospital(approve number Z0553-01). Informed consent was obtained from all women to participate in the study.

## Consent for publication Not applicable

### Funding

This work was supported by the National Natural Science Foundation of China under Grant [number 81300503], and the Province Natural Science Foundation of Hunan, China [number 2023JJ30754]

## Availability of data and materials

The data of this study are available on request from the first author.

## Authors’ contributions

Yan Zhong: Conceptualization, Methodology, Formal analysis, Visualization, Writing – original draft

Li Peng: Resource, Methodology, Formal analysis,Writing – review & editing. Weisi Lai: Resource, Writing – review & editing.

Jian Huang: Conceptualization, Supervision, Writing – review & editing

## Acknowledgements

The authors would like to thank the study staff, laboratory technicians, data teams and co-investigators

## Captions for Table Plus

Table plus 1 Differentially Expressed LncRNAs

Table plus 2 Differentially Expressed mRNAs

Table plus 3 hsa_pathway Result

Table plus 4 Antisense LncRNAs and coding gene data

Table plus 5 LincRNAs associated coding gene data

